# Climatic curves as predictors in MaxEnt niche modeling

**DOI:** 10.1101/2021.11.08.467713

**Authors:** Ondřej Mikula

## Abstract

Environmental niche modelling (ENM) uses different types of variables to predict species occurrence. In widespread use are variables derived from climatic curves, i.e., average annual changes in some climatic parameter. This study shows how to use the climatic curves themselves as ENM predictors. The key step is projection of the curves’ constituent variables on a suitable spline basis, which preserves time-ordering of the variables and supports smoothness of predictions. Complexity of the model is controlled by sensible choice of the spline basis, followed by lasso regularization in model fitting.

## Introduction

Following the seminal paper of Phillips, Anderson & Schapire (2006), the maximum entropy (MaxEnt) modeling has become, arguably, the most widespread method of environmental niche modeling (ENM; a. k. a. species distribution modeling, SDM). Two features of the method make it particularly attractive. First, and most importantly, it deals with presence-only data. This is the far most common type of distribution data, because any observation of the species provides evidence of its presence, while it is quite hard and sometimes impossible to confirm its absence. Second, the method can be made extremely flexible. Due to the use of lasso regularization (Tibshirani 1996) it can deal with thousands of predictors, which allows use of basis transformation approach. A limited set of predictor variables can be transformed into a large number of derived variables that are used as MaxEnt’s actual predictors. This way the model can include even threshold and other nonlinear effects. The original implementation in software *MaxEnt* offers five such transforms, called linear, quadratic, product, threshold and hinge features (Phillips, Anderson & Schapire 2006, Phillips & Dudík 2008). The ‘linear feature’ is in fact an identity transform and makes the software extensible – one can apply any other transform externally, import the result into MaxEnt and leave it unchanged (Merow, Smith & Silander 2013). This approach was made even easier with current implementation in the R package *maxnet* (Phillips et al. 2017), which builds on the package *glmnet* (Friedman, Hastie & Tibshirani 2010) for the regularized fit if generalized linear models. The implementation in R (R Core Team 2021) allows joining customary transforms and MaxEnt fitting into a single reproducible pipeline. The pipeline can further include evaluation of the model using information criteria, e.g., *AICc* (Burnham & Anderson 2002). The MaxEnt model was shown to be an instance of generalized linear model (Fithian & Hastie 2013) and thus it is only natural to use these likelihood-based criteria to compare performance of different variables and their transforms (Warren & Seifert 2001).

The choice of original (non-transformed) variables may reflect prior knowledge of important environmental factors, but no such information is available in many cases and some general set of variables is used, instead. A typical example of the latter is the set of 19 bioclimatic (BIOCLIM) variables, originally defined by Nix (1986), which are increasingly used in ENM as a kind of universal climatic niche descriptors. They include, for instance, ‘average annual temperature’ or ‘precipitation of wettest quarter’, but all nineteen are ultimately transforms of even more basic variables: monthly average precipitations, minimum temperatures and maximum temperatures. At any single place, climate is described by these three climatic curves, each consisting of 12 monthly values. The BIOCLIM variable “ precipitation of wettest quarter”, for instance, is then derived as the maximum summed precipitation in any three consecutive months in the year. Although one could define additional BIOCLIM variables (Kriticos et al. 2014), the final set of predictors is always a constrained version of the original environmental space established by the three curves. MaxEnt fit based on the curves themselves could target variety of other, perhaps unforeseen, climatic features. For instance, a coincidence of steeply decreasing precipitation with steeply increasing temperature can prove predictive for some species. More generally, the use of climatic curves is favorable when the shape of annual trends is important for species distribution.

The aim of the study is to elaborate on this strategy of using climatic curves as predictors in MaxEnt niche modeling. It shows how to treat curves as curves, not as collections of separate monthly values, how to optimize the fit and how to deal with time shifts of otherwise identical local curves. After introducing the methods, we test them on a simulated data set, benchmark data sets of Phillips et al. (2006) and our own distribution data of African subterranean rodents.

## Methods

### Natural cubic spline basis

An example of climatic curves is presented in Figure 1A. Every curve there consists of 12 ordered monthly precipitations in a single cell. The red curves correspond to cells with presence records, while the grey ones to background cells. One could use these 12 variables as an input for MaxEnt modeling, but it would discard all information about their ordering. In fact, it would mean to use not the *curves*, but their elements as the predictors. Only when taking relations between monthly averages into account, one can speak about using the curves. One way to achieve this is to project the variables on a spline basis and use these transformed values as MaxEnt predictors (Hastie, Tibshirani & Friedman 2009 p. 148-150).

**Figure 1.**
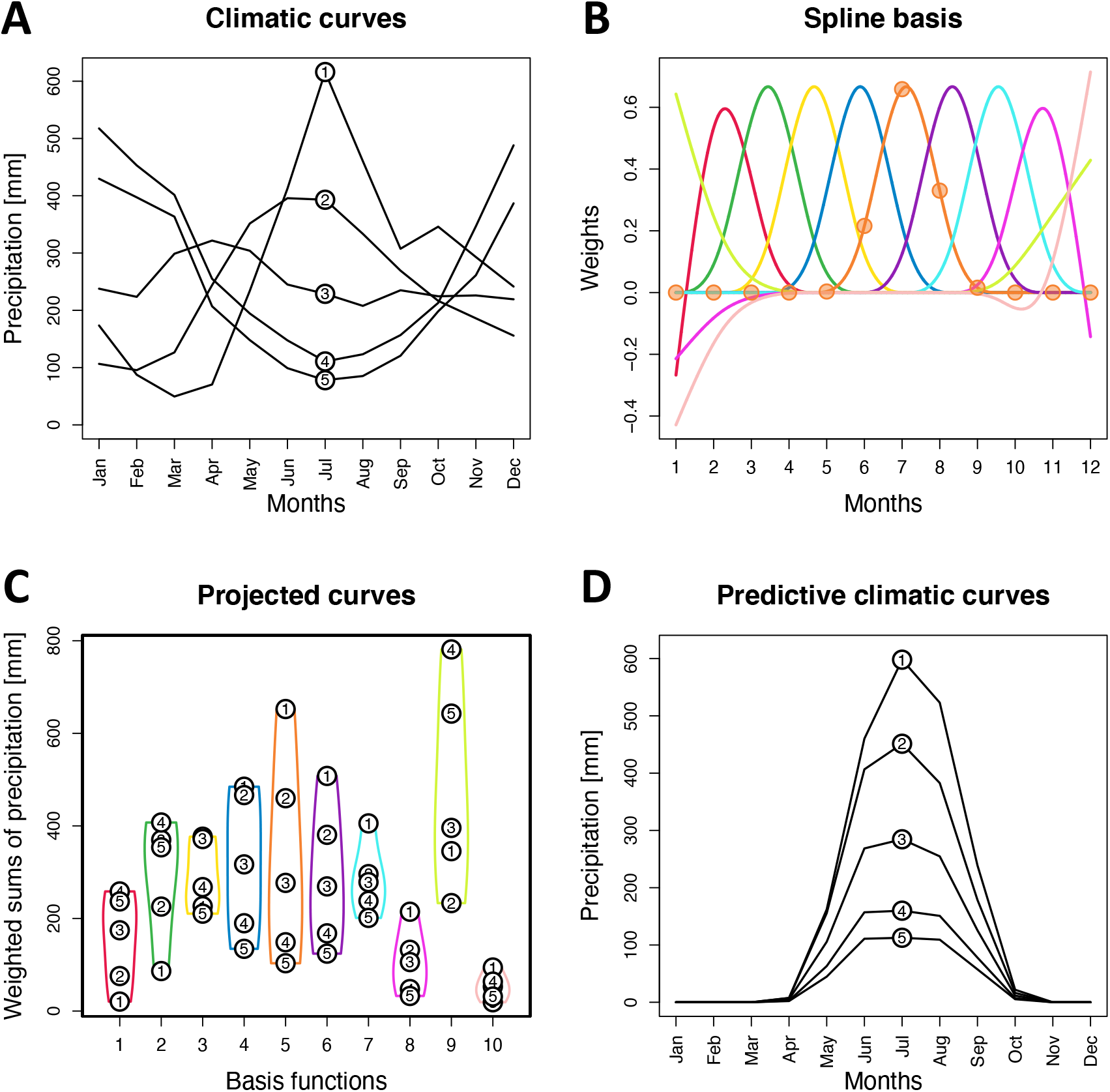
Projection of climatic curves onto the natural cubic spline basis. (A) sample of five presence-point climatic curves; (B) ten spline basis functions (lines) and the values evaluated at ℳ ∈ (1,2, …, 12) for the fifth function (points); (C) curves projected onto the spline basis, i.e., converted into a vector of ten different weighted sums (projections of the five sampled curves are indicated by numbers; violin plots show distribution of transformed values in the underlying data set); (D) predictive features of the five sampled curves under assumption that 4^th^ to 6^th^ basis functions were equally predictive (with MaxEnt regression coefficients *β* = 1) and the remaining function played no role (*β* = 0).

In this paper, we will use a natural cubic spline basis, which is defined by *k* internal knots, i.e., fixed values inside the numeric sequence defining the curve. In our example, the curve-defining numeric sequence is the succession of months and *k* knots can be chosen from the interval (1,12). The natural cubic spline basis is designed to predict a set of piecewise polynomials, which have continuous first and second derivatives at the knots and which are linear beyond the boundary knots. An example of the natural cubic spline basis is given in Figure 1B. This basis has *k* = 8 knots placed equidistantly between boundary points 1 and 12 and consists of *l* = *k* + 2 = 10 functions. These basis functions are in fact latent variables corresponding to different annual trends. When evaluated at values ℳ ∈ (1,2, …, 12), they specify weights given to monthly averages. Calculating these weights is a crucial step, because each basis function is now represented by twelve weights, that can be used to turn a climatic curve into its weighted sum. If a climatic curve is projected onto the basis with 10 functions, 12 monthly averages are replaced by 10 weighted sums (Figure 1C), which can be used as MaxEnt predictors.

In matrix notation, the transform can be written as: **T**=**CB**. Here, **C** is *n* × *m* matrix of climatic curves spanning *m* periods (e.g., twelve months) at *n* sites, **B** is *m* × *l* spline basis with *l* functions evaluated at ℳ ∈ (1,2, …, *m*) and **T** is the matrix of *n* × *l* transformed variables (weighted sums). The backwards projection **C**=**TB′**, where **B′** is the transpose of **B**, restores the original climatic curves up to degree given by complexity of the basis. A basis with more functions can represent the curves in more detail. However, the MaxEnt model gives different weights (regression coefficients) to the transformed variables stored in **T**. These coefficients (in a diagonal matrix **M** of dimension *l*× *l*) can modify the backwards projection (**Ĉ**=**TMB′)**, so it returns re-weighted versions of the climatic curves. These re-weighted curves have those features highlighted, which serve well to predict distribution of the species (Figure 1D).

### Selection of the optimal spline basis

As noted above, the basis with higher *k* can represent the observed curves in more detail. This is not always beneficial, however. Although it can retain minor yet important features, it can also cause overfitting with MaxEnt model based on unimportant particulars of the curves observed in given presence cells. Such model would produce poor predictions in all other cells. This is an example of the usual variance-bias trade-off: predictions of more parameter-rich models are more precise, but it comes at the expense of their accuracy (Anderson 2008, Hastie, Tibshirani & Friedman 2009). One way to optimize model complexity is to minimize Akaike Information Criterion (*AIC*, Akaike 1974). In this paper, we will use *AIC* with second-order bias correction defined as: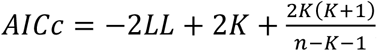, where *LL* is log-likelihood of the model and *K* is the number of estimable parameters (Burnham & Anderson 2002 p. 66). In the model selection procedure spline bases of increasing complexity (=increasing number of basis functions) are used to generate predictors for otherwise identical MaxEnt models and the model with minimum *AICc* is selected as the best one. In addition, *AICc* can be converted to Akaike weights (*w*), which are relative probabilities of models being the best ones (Burnham & Anderson 2002 p. 75).

At the same time, MaxEnt includes yet another regularization of model complexity. While the choice of spline basis determines the number of MaxEnt predictors and the scale of variation they cover, lasso regularization affects how many predictors have non-zero regression coefficients and hence any impact on model predictions. The strength of regularization is given by the lasso penalty multiplier (*λ*) whose value should be set to optimize predictive performance of the model. The optimization of *λ* could also be performed using *AICc*, but in the *maxnet* implementation, it is integral to model fit and it will not be discussed here further.

### Superimposition of climatic curves

The species presence in a cell can be affected by shape of the climatic curve, but not by its position. It possibly matters how much precipitation of the wettest month exceeds the average annual precipitation, but if we do not consider any other variables (like insolation), it does not matter whether the wettest month is January or June. In the methodology described so far, however, precipitations observed in a particular month (e.g., in June) correspond to each other across the whole data set. If precipitation was the only strong predictor of the species presence, it would make much better sense to superimpose the curves prior to spline transformation, i.e., to shift them in time so they fit each other as much as possible. In practice, such pre-processing requires an algorithm exploring possible shifts of cell-specific curves. Here, we suggest an iterative procedure analogous to Generalized Procrustes analysis of Gower (1975).

Initially, an arbitrary curve serves as a template and all other curves are shifted to fit it as much as possible using Euclidean distance as a criterion of the fit quality. Then the average values are calculated for the reordered months, i.e., the average of the first month is the average of the template January and a mixture of months from the other curves, according to their shift. This average curve serves as a template in the next iteration of the procedure, which proceeds until convergence, i.e., when the distance between template and its update becomes negligible (lower than specified threshold).

Using more climatic variables, e.g., precipitation and temperature, makes the superimposition more complex. In this case, each variable has its own template (precipitations are superimposed on precipitations and temperatures on temperatures), but their shifts must be coordinated to preserve local associations of different variables. Assessment of the fit quality on more than one variable requires rescaling all variables to unit variance so their curve-to-template distances are comparable and can be summed up. Moreover, the choice of climate variables driving superimposition is principally detached from that involved in the modelling itself. One can choose just a subset of variables or even a variable not involved in modelling to assess proper superimposition and then shift the curves of other variables accordingly. For instance, one can use precipitation and mean temperature for assessment if superimposition, but precipitation, minimum and maximum temperature for the modelling.

Similarly, one can let just a subset of local climate curves to drive superimposition and then to project the other curves to the final average curve in a single additional superimposition step. An important application of this approach comes with predicting species distribution in the past or in the future. Including climatic curves from the different time layer in the superimposition may distort relative shifts of present curves, which are used in model training, and hence the accuracy of model predictions may be impaired. On the other hand, without superimposition the past or future curves are not comparable to the present ones. It is, therefore, advisable to perform superimposition on the training (i.e., usually the present day) data only and then to project the other curves to the present average curve.

Finally, the superimposition is not an automatic choice, because it can be beneficial as well as detrimental to model performance. There should be either theoretical justification or statistical support for its application. From the model selection point of view original and superimposed curves are different predictors whose performance can be compared, e.g., by information criteria as discussed before.

### Software and data

All analyses presented in this paper were performed in R using the package *maxnet*. The scripts replicating them can be downloaded at …, together with necessary data. The colour scale used in Figure 1 was created by A. Trubetskoy and can be accessed at https://sashamaps.net/docs/resources/20-colors.

All climatic data in this study were downloaded from WorldClim v2 data base (Fick & Hijmans 2017) in 10’ (=1/6°) resolution. This way we obtained data for 90563cells covering the continent of Africa (for analysis of simulated and empirical data sets) and for 57913 cells covering South America and Carribean (for analysis of benchmark data sets). Climatic curves were assembled from WorldClim raster layers of monthly average precipitation, minimum temperatures and maximum temperatures, 19 BIOCLIM variables were downloaded separately in the form of pre-prepared raster layers. Estimates of the last glacial maximum climate were produced by the community climate system model v. 4 (Gent et al. 2011) and accessed from WorldClim v1 (Hijmans et al. 2005).

## Results

*to be done*

## Discussion

*to be done*

